# Tau-related reduction of glucose metabolism in mild cognitive impairment occurs independently of APOE ε4 genotype and is gradually modulated by β-amyloid

**DOI:** 10.1101/2024.01.23.576866

**Authors:** Felix Carbonell, Carolann McNicoll, Alex P. Zijdenbos, Barry J. Bedell, Alzheimer’s Disease Neuroimaging Initiative

## Abstract

**Background:** PET imaging studies have shown that spatially distributed measurements of β-amyloid are significantly correlated with glucose metabolism in Mild Cognitive Impairment (MCI) independently of the APOE ε4 genotype. In contrast, the relationship between tau and glucose metabolism at different stages of Alzheimer’s Disease (AD) has not been fully understood.

**Objective:** We hypothesize that spatially distributed scores of tau PET are associated with an even stronger reduction of glucose metabolism, independent of the APOE ε4 genotype and gradually modulated by β-amyloid.

**Methods:** We applied a cross-sectional statistical analysis to concurrent [18F]flortaucipir PET, [18F]florbetapir PET, and 2-[18F]fluoro-2-deoxyglucose (FDG) PET images from the Alzheimer’s Disease Neuroimaging Initiative (ADNI) study. We employed a Singular Value Decomposition (SVD) approach to the cross-correlation matrix between tau and the FDG images, as well as between tau and β-amyloid PET images. The resulting SVD-based tau scores are associated with cortical regions where a reduced glucose metabolism is maximally correlated with distributed patterns of tau, accounting for the effect of spatially distributed β-amyloid.

**Results:** From a population of MCI subjects, we found that the SVD-based tau scores had their maximal spatial representation within the entorhinal cortex and the lateral inferior temporal gyrus, and were significantly correlated with glucose metabolism in several cortical regions, independently from the confounding effect of the β-amyloid scores and APOE ε4. Moreover, β-amyloid gradually modulated the association between tau and glucose metabolism.

**Conclusions:** Our approach uncovered spatially distributed patterns of the tau-glucose metabolism relationship after accounting for the β-amyloid effects. We showed that the SVD-based tau scores have a strong relationship with decreasing glucose metabolism. By highlighting the more significant role of tau, rather than β-amyloid, on the reduction of glucose metabolism, our results could have important consequences in the therapeutic treatment of AD.

## Introduction

Preliminary empirical evidence about the pathogenesis of Alzheimer’s disease (AD) led to the formulation of the so-called “amyloid cascade hypothesis”[1–3]. In general terms, the amyloid cascade hypothesis posits that the deposition and widespread accumulation of β-amyloid are the key triggering processes inducing a cascade of time-ordered events, including tau pathology, glucose hypometabolism, brain atrophy, and eventually, cognitive impairment and dementia. In contrast, more recent studies validate the alternative hypothesis that not only the deposition of β-amyloid plaques, but also the accumulation of neurofibrillary tangles of tau aggregates acting in a coordinated fashion, lead to the ultimate neuronal loss and cognitive deterioration [4–6]. As such, investigators have made substantial efforts to gain a better understanding of the underlying relationships between β-amyloid, tau, and glucose hypometabolism, namely, the main AD-associated pathophysiological biomarkers involved in the amyloid cascade hypothesis [7–11]. A better understanding of the role played by tau, β-amyloid, and their synergistic interaction in neurodegeneration across different AD stages would also provide more solid foundations for effective therapeutic treatments and drug development to prevent and treat AD. Indeed, if β-amyloid has little or no influence on neurodegeneration markers of AD, it would be preferable to redirect efforts towards the development of therapeutics that target tau at pre-symptomatic stages of the disease. In contrast, if a synergistic interaction between tau and β-amyloid is the primary driver of neurodegeneration, then therapeutic agents targeting tau and β-amyloid concomitantly would be an appropriate path to follow.

Extensive studies have been conducted to elucidate the impact of β-amyloid and APOE ε4 genotype on the reduction of glucose metabolism during the mild cognitive impairment (MCI) stage of AD [12–19]. While it was initially reported that APOE ε4, and not global measures of β-amyloid, contribute to glucose hypometabolism in cognitively normal (CN) according to Jagust *et al.* [15] and MCI subjects as per Carbonell *et al.* [13], we recently showed [14] that spatially distributed β-amyloid scores produce a stronger relationship with decreasing glucose metabolism than APOE ε4 genotype and global measures of β-amyloid. This apparent contradiction came from the fact that such initial reports employed a representation of β-amyloid by a single global index with little or no spatial specificity, while our more recent study [14] used a whole-brain Singular Value Decomposition (SVD) approach from Worsley *et al.* [20] to derive a more accurate representation of β-amyloid scores that were intended to be maximally correlated with glucose metabolism. Hence, in concordance with previous studies by Lowe *et al.* [17], our more recent results on this subject [14] reinforce the idea that, independently of the APOE ε4 genotype, β-amyloid accumulation and reduction of glucose metabolism are more likely to occur simultaneously throughout the clinical spectrum of AD progression in a β-amyloid-dependent manner. Thus, an accepted consensus [13,16,17,21] is that even though the APOE ε4 genotype alone can be related to hypometabolism, when interacting with β-amyloid deposition, most of the reduction in metabolism is attributable to the latter rather than the former. However, as pointed out in Knopman *et al.*, Lowe *et al.*, and Mormino *et al.* [16,17,21], such interpretation should not be taken as APOE ε4 genotype and amyloid-β burden providing redundant information, but as having an additive impact on the reduction of glucose metabolism.

Much less consensus has been reached about the association between tau accumulation and the reduction of glucose metabolism, as well as its impact on the cognitive deterioration that eventually leads to dementia. Preliminary studies argued that tau could be simply considered a mediator between β-amyloid and neurodegeneration [22–25], despite some other results suggesting a more complex interpretation and pointing to a synergistic interaction between neocortical tau and β-amyloid in relation to glucose metabolism [26,27]. In turn, AD-related processes, such as reduction of glucose metabolism or gray matter volume loss, have also been thought to play a mediating role between tau uptake and cognition deterioration [28–30].

Nevertheless, the number of contradictory results emerging from the previous studies is the main reason impeding a consensus about the actual molecular mechanisms that relate glucose metabolism, β-amyloid deposition, and tau accumulation to AD progression.

The first *in vivo* evidence of the positive association between tau uptake and glucose hypometabolism was provided by Bischof *et al.* [26]. Although employing a small sample size of AD subjects, it was also shown that spatially concurrent (i.e., same spatial location) β-amyloid burden and tau deposition had an interactive effect on glucose metabolism [26]. In particular, the increase of β-amyloid was associated with a stronger impact of tau accumulation on the reduction of glucose metabolism [26]. By using a larger sample of typical and atypical AD subjects with high levels of β-amyloid, Whitwell *et al.* [23] have shown that regional hypometabolism was more strongly correlated to regional tau pathology than to regional β-amyloid burden. Consistent with these early results, Strom *et al.* [31] more recently demonstrated that, independently of the APOE ε4 genotype effect, local tau pathology was associated with local hypometabolism in a sample of symptomatic AD subjects.

Similarly, using a large sample of CN older adults, Hanseeuw *et al.* [27] showed that the increase in local tau pathology was associated with local hypometabolism, particularly within the entorhinal cortex and the inferior temporal cortices. Additionally, by using a network-based perspective, they demonstrated that tau tracer uptake in the entorhinal cortex was not only related to local hypometabolism, but extended to nearby temporal regions and the cingulate cortex [27]. Hanseeuw *et al.* [27] also showed that the local negative correlation between tau and FDG increased in those subjects with high levels of global β-amyloid burden, taken from [11C]PIB PET measurements of an AD-signature composite ROI. In contrast, they reported that in those participants with low levels of β-amyloid, the increase of tau signal in the inferior temporal regions was associated with increased glucose metabolism [27]. This surprising amyloid-dependent opposing association between tau and glucose metabolism has been further corroborated by Adams *et al.* [32] using a large-scale sparse canonical correlation approach in a sample of CN subjects. Indeed, the analysis carried out by Adams *et al.* in [32] showed that, within a regime of low levels of β-amyloid, overall increases in tau tracer binding were mostly associated with increases in glucose metabolism, particularly in the frontal and medial temporal lobes. In contrast, at high levels of β-amyloid, correlations between tau and FDG were predominantly negative all over the brain [32]. The same sort of contrasting amyloid-dependent associations between FDG PET and tau PET have been replicated in a sample of MCI subjects by Rubinski *et al.* [33]. For low levels of global β-amyloid, increases in tau signal were locally associated with increasing glucose metabolism in several regions, including the parietal lobes and cingulate cortices [33]. In contrast, negative associations between FDG signal and tau were only observed for the positive β-amyloid cases [33].

The number of contrasting results suggests that the association between tau and glucose metabolism in CN and MCI depends on the levels of β-amyloid in a spatially distinct manner. However, previous studies involved a series of methodological constraints that have precluded a better understanding of their associated biological findings. Indeed, with a few exceptions from Adams *et al.* [32], most of the correlational analysis between tau PET and FDG PET have been carried out at a locally spatial level and/or employing a single global measure for the quantification of the β-amyloid accumulation. This approach is a clear limitation given that some previous studies have remarked the importance of considering large-scale multivariate techniques between PET images to uncover hidden patterns of associations between β-amyloid and FDG in Carbonell *et al.* [14], as well as between β-amyloid and tau in Carbonell *et al.* [34]. In this work, we not only investigate the effect of tau accumulation, but also examine a potential synergy between tau, β-amyloid, and the APOE ε4 genotype on the reduction of glucose metabolism in subjects with MCI. Given previous limitations regarding the spatial scale, we employ a SVD approach to produce data-driven scores that reveal whole-brain patterns in the cross-correlation structure between FDG PET and tau PET. Thus, this work extends our previous SVD cross-correlation analysis [14], which uncovered hidden patterns of the glucose metabolism-amyloid-β relationship. We hypothesize that SVD-based tau scores are associated with a stronger reduction of glucose metabolism independently of the effects of the APOE ε4 genotype. We also hypothesize that the spatial association between tau and glucose metabolism not only depends on the global amyloid status, but varies gradually according to the spatially distributed measures of β-amyloid.

## Materials and Methods

### Subjects and Image Acquisition

Data used in the preparation of this article were obtained from the ADNI database (http://adni.loni.usc.edu). The ADNI was launched in 2003 by the National Institute on Aging (NIA), the National Institute of Biomedical Imaging and Bioengineering (NIBIB), the Food and Drug Administration (FDA), private pharmaceutical companies, and non-profit organizations, as a $60 million, 5-year public-private partnership, which has since been extended. ADNI is the result of the efforts of many co-investigators from a broad range of academic institutions and private corporations, and subjects have been recruited from over 55 sites across the U.S. and Canada. To date, the ADNI, AND-GO, ADNI-2 and ADNI-3 protocols have recruited over 1,500 adults, ages 55 to 90, to participate in the research, consisting of cognitively normal (CN) older individuals, people with early or late Mild Cognitive Impairment (MCI), and people with early AD. For up-to-date information, see www.adni-info.org.

The subjects of this cross-sectional study consisted of 119 subjects, ranging from Early to Late MCI, from the ADNI study who had available 2-[18F]fluoro-2-deoxyglucose (FDG), [18F]flortaucipir PET, and 3D T1-weighted anatomical MRI within a time frame of 6 months. Early MCI (EMCI) subjects had MMSE scores between 24 and 30 inclusively, a CDR of 0.5, a reported subjective memory concern, an absence of dementia, an objective memory loss measured by education-adjusted scores on delayed recall of one paragraph from the Wechsler Memory Scale Logical Memory (WMSLM) II, essentially preserved activities of daily living, and no impairment in other cognitive domains. Late MCI subjects had the same inclusion criteria, except for objective memory loss measured by education-adjusted scores on delayed recall of one paragraph from (WMSLM) II.

A detailed description of the ADNI MRI and PET image acquisition protocols can be found at http://adni.loni.usc.edu/methods. ADNI studies are conducted in accordance with the Good Clinical Practice guidelines, the Declaration of Helsinki, and U.S. 21 CFR Part 50 (Protection of Human Subjects) and Part 56 (Institutional Review Boards), where informed written consent was obtained from all participants at each site.

Out of the whole sample of 119 subjects, 116 subjects also had concurrent [18F]florbetapir PET scans within a time frame of 6 months. Those 116 [18F]florbetapir PET scans were classified into high (Aβ_H_) and low (Aβ_L_) amyloid subjects according to their Standardized Uptake Value Ratio (SUVR) values in an optimal target ROI (Stat-ROI) including areas of the posterior cingulate cortex, precuneus, and medial frontal cortex, with an associated cutoff of SUVR = 1.24. That optimal target ROI and the cutoff value for the segregation of subjects into low and high amyloid were selected by our data-driven approach that has been previously developed by Carbonell *et al.* [35].

### Image Processing

MR and PET images were processed using the PIANO™ software package (Biospective Inc., Montreal, Canada). T1-weighted MRI volumes underwent image non-uniformity correction using the N3 algorithm from Sled *et al.* [36], brain masking, linear spatial normalization utilizing a 9-parameter affine transformation, and nonlinear spatial normalization to map individual images from native coordinate space to Montreal Neurological Institute (MNI) reference space using a customized, anatomical MRI template derived from ADNI subjects. The resulting image volumes were segmented into gray matter (GM), white matter (WM), and cerebrospinal fluid (CSF) using an artificial neural network classifier from Zijdenbos *et al.* [37] and partial volume estimation from Tohka *et al.* [38].

The [18F]FDG, [18F]florbetapir, and [18F]flortaucipir PET images underwent several pre-processing steps, including frame-to-frame linear motion correction, smoothing with scanner-specific blurring kernels to achieve 8mm FWHM per Carbonell *et al.* [39], and averaging of dynamic frames into a static image. The resulting smoothed PET volumes were linearly registered to the subject’s T1-weighted MRI and, subsequently, spatially normalized to reference space using the linear and nonlinear transformations derived from the anatomical MRI registration. The GM density map for each subject was transformed to the same final spatial resolution (i.e., re-sampled to the same voxel size) as the PET data in order to account for the confounding effects of atrophy in the group-level statistical model. SUVR maps of the PET images were generated from [18F]florbetapir and [18F]flortaucipir PET using the gray matter-masked full cerebellum as a reference region. The FDG SUVR maps were generated with the pons as a reference region.

To minimize PET off-target binding in non-gray matter regions, we have restricted our analysis to those voxels in the brain cortex that have been previously masked by a GM mask in the nonlinear template space. Hence, all our voxelwise PET maps and statistical maps were projected onto the cortical surface for visualization purposes only.

### Statistical Analysis

The main component of the methodology employed here is the Singular Value Decomposition (SVD) of the multi-modality cross-correlation matrix between the tau PET and FDG PET images demonstrated by Carbonell *et al.* [14]. Briefly, the SVD of the cross-correlation matrix *C* between the tau and the FDG dataset can be expressed as *C* = *UWV*’, where *U* and *V* are orthonormal matrices, whose columns are the so-called eigenimages or spatial loadings, and *W* is a diagonal matrix of ordered eigenvalues (see details in Carbonell *et al.* [14]). For practical reasons and an easier interpretation, *C* is typically approximated by the first few components, ordered according to the values of the weights in *W*. Hence, the cross-product of the first spatial loadings would produce the largest additive component of the full voxels x voxels matrix *C,* without the need for reconstructing it or performing the cross-product operation for extracting significant information about the underlying cross-correlation patterns.

Correspondingly, the SVD analysis also provides individual scores for each PET modality that can be computed by simply projecting the spatial loadings onto the corresponding PET dataset. Thus, for a more precise statistical inference, those individual scores can be included as regressors in a voxelwise General Linear Model (GLM). In particular, we used GLM for statistical assessment of the effect of SVD-based tau scores on FDG PET SUVR. As previously explained by Carbonell *et al.* [14], we employed a leave-one-out cross-validation technique to produce the individual SVD-based scores. Thus, leaving each sample out one-at-a-time, the SVD and corresponding eigenimages are produced from the rest of the sample. Then, the individual scores of SUVR_SVD_ for the left-out sample are computed by mapping them onto the space of the orthogonal eigenimages corresponding to the rest of the sample. Such a cross-validation approach overcomes any possible circularity effect produced by the computation of the SVD components and their subsequent inclusion into a GLM.

Several models were assessed in our analysis. The first model was intended to evaluate the effect of the SVD-based tau score and APOE ε4 status on FDG signal:

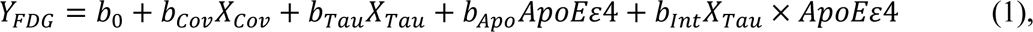

where *Y_FDG_* denotes the FDG SUVR as a predicting variable; *X*_Cov_ includes age, sex, and cognitive measurements (MMSE and ADAS-Cog) as global confounding covariates; GM density as a voxelwise confounding covariate; and SVD-based tau scores (*X*_Tau_), APOE ε4status, and tau x APOE ε4 status interaction as predictors of interest.

A second model was assessed to evaluate the interactive effect of β-amyloid and tau on FDG:

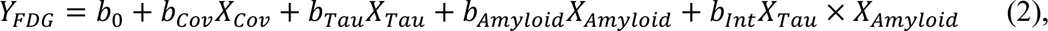

where, in this case, *X*_Cov_ also includes the APOE ε4 status and *X*_Amyloid_ denotes SVD-based amyloid scores resulting from an independent SVD analysis between FDG and amyloid PET images. Notice also that the previous model can be rewritten as:

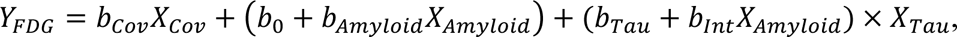

which can be used to assess how the relationship between FDG and tau can be continuously modulated by amyloid. Therefore, when there is a true interaction (e.g., assessed by *b*_lnt_) between *X*_Amyloid_ and *X*_Tau_, it can be inferred that the main effect of tau over FDG (i.e., *b*_Tau_ + *b*_lnt_*X*_Amyloid_) depends on, or is conditional on, the amyloid value *X*_Amyloid_. Similar to our previous metabolic connectivity analysis by Carbonell *et al.* [12], we estimate the strength of the association between FDG and tau any particular value *s* of amyloid by using the t-test associated with the “slope” *b*_Tau_ + *b*_lnt_*X*_Amyloid_ at *X*_Amyloid_ = *s*:

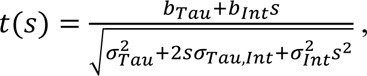

where σ_Tau_, σ_lnt_ denote the standard deviation of *b*_Tau_ and *b*_lnt_, respectively, and σ_Tau,lnt_ denotes the covariance between *b*_Tau_ and *b*_lnt_.

A third model included local tau measurements (*XSeed*_Tau_) taken from 5 mm seeds centered on areas highly contributing (e.g., local maxima) to the first SVD-based tau eigenimage:

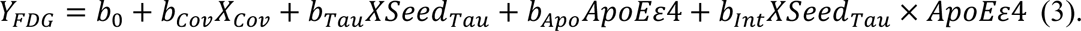

Our intention here was to reveal local-to-distributed patterns on the relationship between (local at the seed level) tau and (distributed) glucose metabolism, as compared to the distributed tau measurements produced by the SVD analysis.

Post-hoc, two-tailed Student’s t-tests were performed to assess the main effects of interest and interaction terms in each of the previous models. The voxelwise statistical analysis was performed using an in-house Python version of the SurfStat toolbox (http://www.math.mcgill.ca/keith/surfstat), where statistical maps were projected onto the cortical surface for visualization purposes only. The t-statistic maps corresponding to each effect of interest were thresholded using the False Discovery Rate (FDR) procedure (α = 0.05) from Genovese *et al.* [40] to control for multiple comparisons.

## Results

The subject characteristics analysis from Table 1 revealed no statistically significant associations between APOE ε4 status and sex (*χ^2^* = 2.59, *p* = 0.11) or age (*t* = 0.26, *p* = 0.63). Neither the MMSE (*t* = 0.23, *p* = 0.72) nor the ADAS-Cog (*t* = 0.89, *p* = 0.07) showed statistically significant differences between the APOE ε4 statuses. A contingency table analysis within the sample of 116 subjects with FDG, tau, and amyloid PET revealed statistically significant associations between APOE ε4 status and amyloid status (*χ^2^* = 8.61, *p* = 0.003). For the sake of this section, any reference to amyloid data will be understood as an analysis restricted to the sample of 116 subjects with FDG, tau, and amyloid PET images.

**Table 1.** Summary of subject characteristics.

|  | All | APOE ε4 Carrier | APOE ε4 Non-Carrier |
| --- | --- | --- | --- |
| **Sample Size** | 119 | 21 | 98 |
| **Age** | 72.32 ± 7.81 | 71.91 ± 8.08 | 72.41 ± 7.79 |
| **Sex (F/M)** | 52/67 | 13/8 | 39/59 |
| **Aβ (Low / High)** | 50/66 | 1/19 | 49/47 |
| **MMSE** | 28.03 ± 1.79 | 27.95 ± 1.74 | 28.05 ± 1.81 |
| **ADAS-Cog** | 12.66 ± 3.96 | 11.97 ± 3.20 | 12.83 ± 4.12 |

The first, second, and third SVD components accounted for 21.49%, 8.81%, and 7.21% of the total co-variability between the FDG and flortaucipir PET images, respectively. Figures 1A and 1B show the spatial representation of the corresponding spatial loadings for each PET modality. The strongest positive weights in the first tau eigenimage (Figure 1A) correspond to the entorhinal cortex, the lateral inferior temporal gyri, the fusiform gyri, the bilateral angular gyrus, as well as areas of the posterior cingulate cortex. Thus, such regions appeared to be negatively correlated to the strongest negative loads in the FDG eigenimage (Figure 1B), which correspond to the lateral inferior temporal gyri, angular gyri, and small areas within the posterior cingulate cortex. In a similar manner, Supplementary Figure 1 shows the spatial loadings for tau and amyloid corresponding to the second and third SVD components. Notice that the second component (Supplementary Figures 1A and 1B) highlights the situation where an increase in tau uptake within areas of the lateral and medial frontal lobules is associated with both a reduction of glucose metabolism in the lateral inferior temporal gyri and an increase of glucose metabolism within the medial fronto-parietal cortex. The third SVD component reflects (Supplementary Figures 1C and 1D) that the increase of tau tracer binding within the medial orbito-frontal cortex and the parahippocampal gyri is associated with an increase of glucose metabolism within the temporo-occipital cortex. Figure 1C shows boxplots corresponding to the individual tau SVD scores for the two classification groups according to both the APOE ε4 status and the amyloid status. The mean values for the tau SVD scores were SUVR = 1.84 ± 0.38, 1.68 ± 0.38, 1.50 ± 0.24, and 1.86 ± 0.41 for the APOE ε4 Carrier, APOE ε4 non-Carrier, Aβ_L_, and Aβ_H_ groups, respectively. Those values resulted in no statistically significant differences between APOE ε4 Carrier and Non-Carrier (*t* = 1.72, *p* = 0.08), but highly significant differences between amyloid Low and High groups (*t* = −5.54, *p* < 0.001). Figure 1D shows the scatter plot for the first tau and FDG SVD-based scores, which produced a large correlation value of *r* = −0.49 (*p*<0.001), providing evidence for an overall linear dependency between FDG and tau. Also notice that, when segregated by β-amyloid, the tau and FDG scores remain strongly negatively correlated.

**Figure 1.**
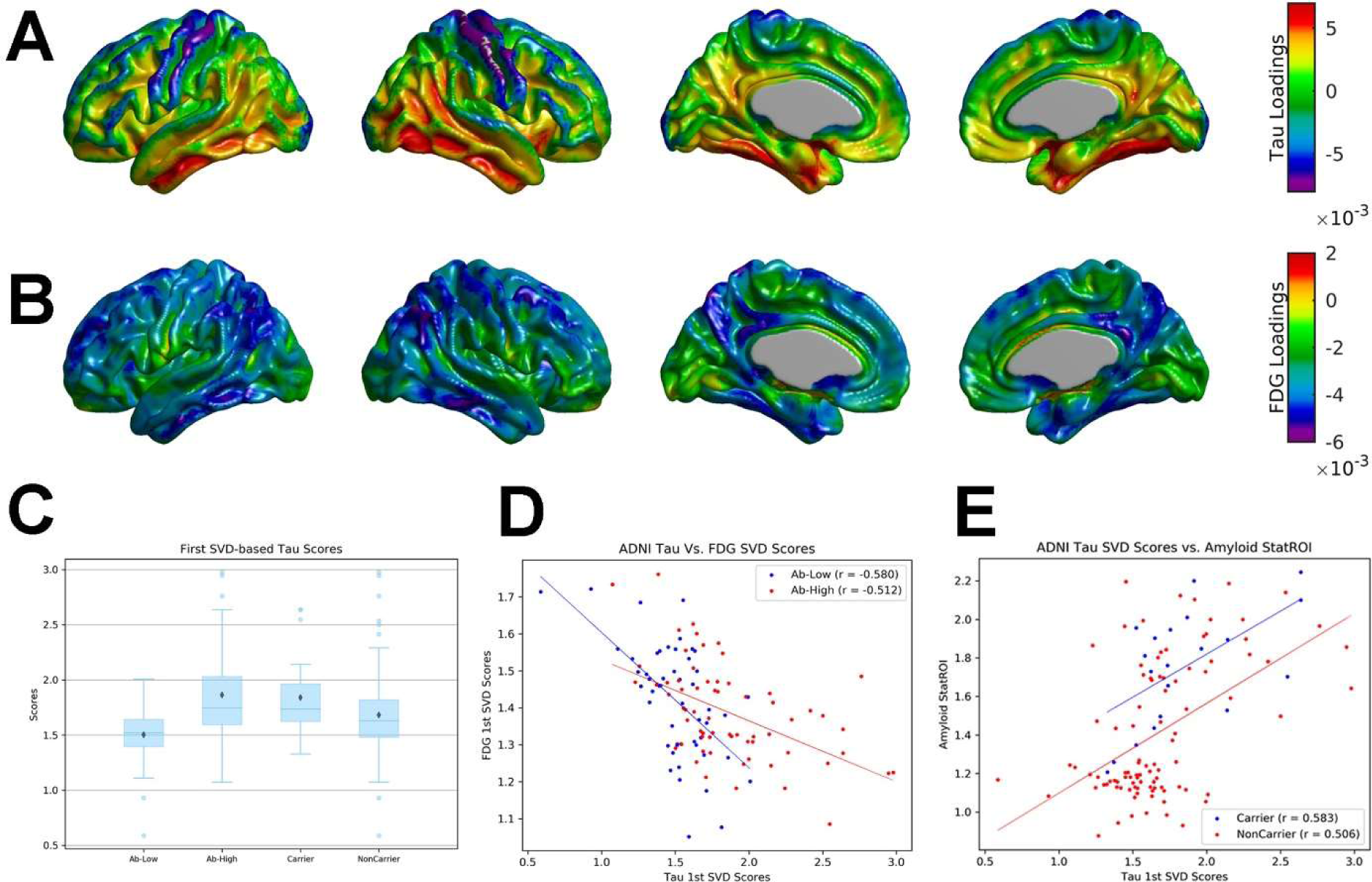
The first SVD component from the cross-correlation between FDG and tau PET images from MCI subjects accounts for 21.49% of the total co-variability. (A) The strongest positive weights in the first tau eigenimage correspond to the medial and lateral inferior temporal gyri. (B) The strongest negative loads in the FDG eigenimage correspond to the lateral inferior temporal gyri, angular gyri, and small areas within the posterior cingulate cortex. (C) Boxplots for the segregation of the tau scores according to APOE ε4 and global β-amyloid statuses shows strong statistically significant differences between amyloid Low and High groups. (D) The scatter plot for the first tau and FDG scores shows large overall and by β-amyloid group correlation values. (E) The strong correlation between the tau scores and the Stat-ROI amyloid measurements does not change with APOE ε4 status.

Similarly, Figure 1E shows that the first tau scores were also significantly correlated to the Stat-ROI measurements (*r =* 0.53, *p <* 0.001). The strength and direction of this linear association did not change when the data was segregated by the APOE ε4 status.

The assessment of the main effects of APOE ε4 status and SVD-based tau scores on FDG SUVR was carried out by statistical inference over the coefficients *b*_Apo_ and *b*_Tau_ in Model (1), respectively. Although showing an overall trend of relationship with glucose metabolism, the main effect of APOE ε4 did not produce areas of statistical significance after controlling for multiple comparisons (Figure 2A). In contrast, the increase in tau scores was significantly associated with a reduction in glucose metabolism in several regions, including the precuneus, the posterior cingulate cortex, the lateral inferior temporal areas, the entorhinal cortex, and extended regions of the parietal cortex (Figure 2B). For comparison purposes, Model (1) was also evaluated to assess the association between glucose metabolism and the SVD-based amyloid scores (i.e., replaced *X*_Tau_ by the SVD-based amyloid scores *X*_Amyloid_). There, only small areas of statistical significance could be observed within the lateral inferior temporal gyri, entorhinal cortex, and the parietal cortex (Figure 2C).

**Figure 2.**
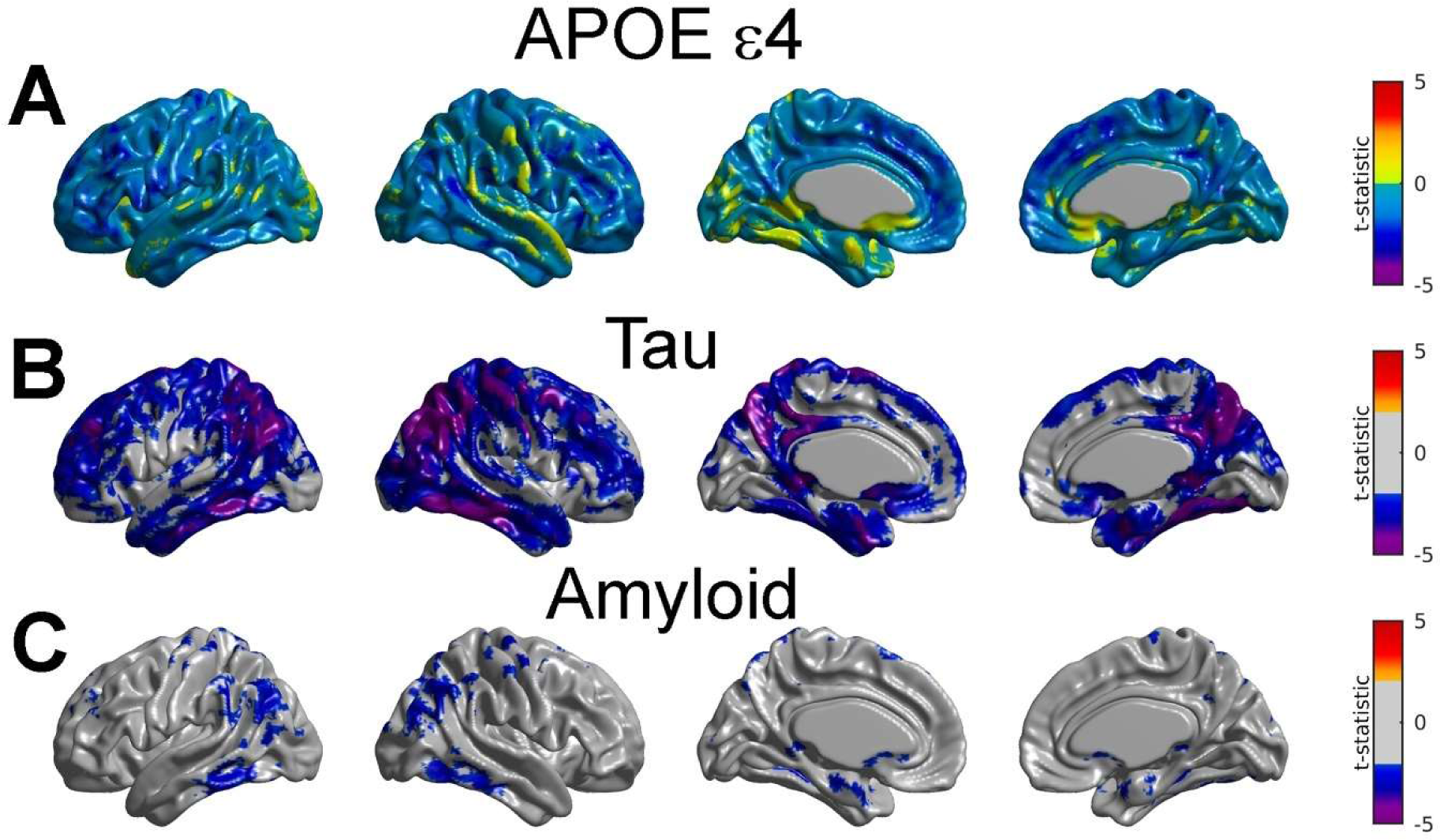
Statistical assessment of APOE ε4, tau scores and β-amyloid scores on FDG SUVR. (A) No region of statistically significant effect APOE ε4 after multiple comparisons FDR thresholding. (B) Tau scores were significantly associated with a reduction in glucose metabolism in several regions, including the precuneus, the posterior cingulate cortex, the lateral inferior temporal areas, the entorhinal cortex, and extended regions of the parietal cortex. (C) β-amyloid scores were significantly associated with FDG in small areas the lateral inferior temporal gyri, entorhinal cortex, and the parietal cortex.

We subsequently evaluated Model (2) to assess the interactive effects of distributed β-amyloid and tau on glucose metabolism. Figure 3A shows that the SVD-based tau scores still produced extended areas of association with the reduction of glucose metabolism after adjusting for the effect of distributed amyloid. In contrast, the main effect of β-amyloid did not produce areas of statistical significance on FDG SUVR (Figure 3B) after adjusting for the effect of distributed tau. Similarly, the interaction term between β-amyloid and tau did not show regions surviving the multiple comparisons thresholding (Figure 3C), although the tau-related reduction of glucose metabolism in areas such as lateral inferior temporal cortex, precuneus, and parietal cortex seems to be relatively associated with an increase of β-amyloid. Notice that, although not statistically significant, the increase in β-amyloid seems to be associated with a positive relationship between FDG and tau in large areas of the cortex, particularly within the frontal cortex (Figure 3C).

**Figure 3.**
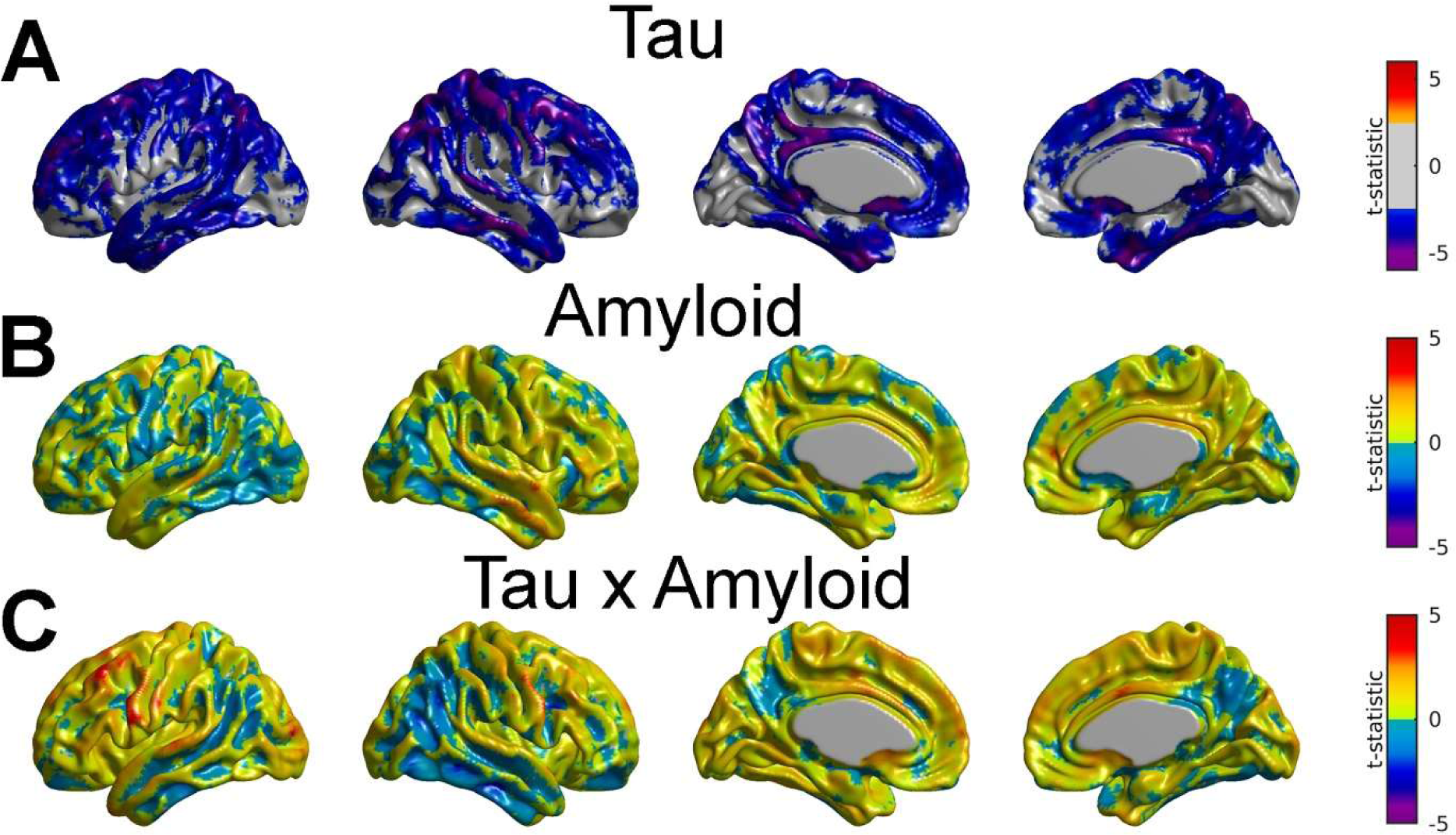
Assessment of the interaction *Amyloid x Tau* effect on FDG PET. (A) The SVD-based tau scores show large areas of association with the reduction of glucose metabolism after adjusting for the effect of β-amyloid. (B) Amyloid did not produce areas of statistical significance on SUVR FDG after adjusting for tau. (C) Interaction between β-amyloid and tau did not show regions surviving the multiple comparisons thresholding.

To remove any potential bias on the choice of the β-amyloid measurements, we repeated the previous analysis by using alternative amyloid scores resulting from the SVD analysis between amyloid and tau PET images (as opposed to amyloid and FDG PET images). The result of such an analysis is shown in Supplementary Figure 2. In this figure, one can still observe extended areas of reduction in glucose metabolism associated with tau scores after controlling for the effects of the alternative β-amyloid scores (the one from the tau-amyloid SVD analysis). Similarly, neither the main effect of β-amyloid nor the interaction between β-amyloid and tau produced statistically significant results.

Figure 4 shows how the relationship between glucose metabolism and the SVD-based tau scores is continuously modulated by the spatially distributed β-amyloid scores. There, one can easily observe an evident overall tau-related reduction in glucose metabolism associated with the increase in β-amyloid levels. For low levels of β-amyloid (Figure 4A), the areas showing the stronger negative tau-FDG correlations correspond to the lateral inferior temporal gyri, angular gyri, precuneus, and posterior cingulate cortex. Notice also small areas of increase in glucose metabolism, particularly in the pars opercularis. With the increase of β-amyloid from SUVR=1.1. to SUVR=1.7, there is an evident increase in the strength of the negative correlations between FDG and tau, as well as a decrease in extent and strength in those small areas associated to a tau-related increase in glucose metabolism (Figures 4B, 4C and 4D). In fact, for very high values of β-amyloid (Figure 4D), the spatial extent and strength of such negative correlations are noticeable for most of the cortex. To further corroborate this modulating effect of β-amyloid, we performed additional SVD analysis between tau and FDG for the two subsets of subjects resulting from segregating the 166 subjects with amyloid PET data into the Aβ_H_ and Aβ_L_ classes. Supplementary Figure 3 shows the corresponding spatial loadings for the first principal component of tau and FDG. A noticeable difference is that the tau loadings appear to be stronger for the Aβ_H_ case, particularly for the lateral inferior temporal gyri and the fusiform gyri. Also, the negative FDG loadings are only strong for the Aβ_H_ case, which provides evidence of the high impact of elevated β-amyloid in the tau-FDG association.

**Figure 4.**
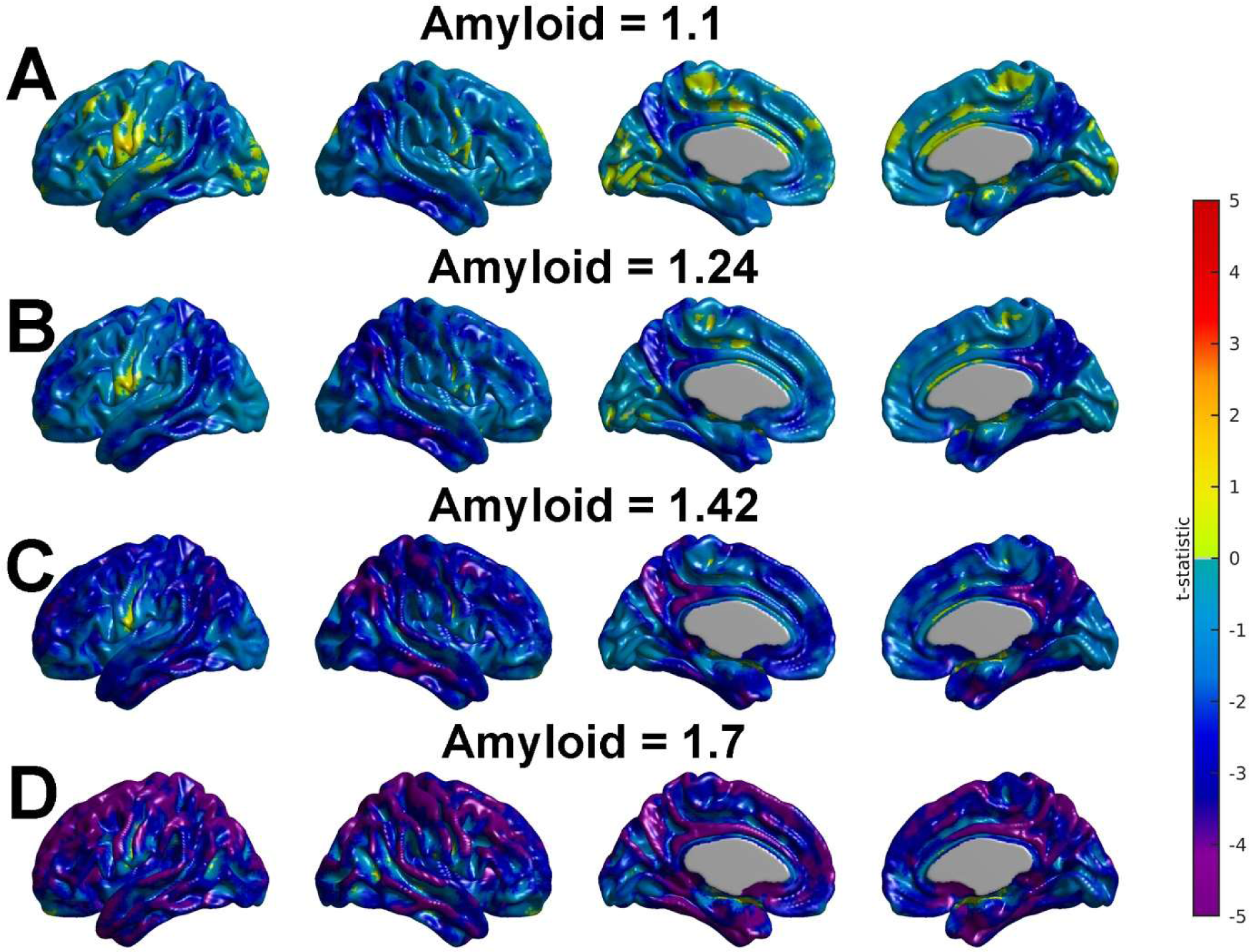
Modulation of the tau-FDG association by β-amyloid. (A) Overall tau-related reduction in glucose metabolism at low level of β-amyloid, particularly within the lateral inferior temporal gyri, angular gyri, precuneus, and posterior cingulate cortex. Small areas of tau-related increase in glucose metabolism, particularly in the pars opercularis. (B, C, D). Gradual increase in the strength and spatial extent of the areas where tau is associated with FDG as β-amyloid increases from SUVR=1.1. to SUVR=1.7.

Several bilateral seed regions were identified as local maxima in the first tau eigenimage: entorhinal cortex (with Talairach coordinates [−26 −7 −29] and [26 −7 −29]), lateral inferior temporal gyri ([−52 −52 −12] and [52 −52 −12]), dorsal posterior cingulate cortex ([−7 −51 27] and [7 −51 27]), fusiform gyri ([−31 −42 −14] and [31 −42 −14]), or local minima in the first FDG image: angular gyri ([−49 −57 35] and [49 −57 35]). A detailed seed-based correlation analysis is presented for two seed regions:(a) the left fusiform gyrus (FUSI) and (b) the right angular gyrus (RANG), which were regions that showed distinct amyloid-glucose metabolism correlation patterns in our previous study [14].

Figures 5A and 5B show the spatial distribution of the tau pattern of spread, or “tau network” associated with the left fusiform and the right angular seeds, respectively. As expected, due to the mathematical properties of Pearson’s correlation coefficient, strong correlation values were obtained around the seed location. Notice that the seed-based tau network relative to the LFUSI seed (Figure 5A) appears to be strong and spatially extended beyond the fusiform gyrus and covers large bilateral areas of the temporal-parietal lobes and precuneus. In contrast, the RANG tau network is particularly strong within the right angular gyrus itself and within the bilateral precuneus and posterior cingulate cortex. Pearson’s correlation between local tau and local glucose metabolism (i.e., at the same seed location) in the LFUSI and RANG were r = −0.44 (p < 0.001) and r = −0.11 (p = 0.28), respectively. Table 2 shows the average tau and FDG SUVR values for the 10 seed regions as well as their (local) correlation values. The RANG and LFUSI tau measurements do not only show a distinctive local correlation pattern with glucose metabolism, but also a distinct local-to-distributed cross-correlation pattern as well. Indeed, Figures 5C and 5D show the t-statistics maps resulting from fitting the Model (3) with LFUSI and RANG seed tau measurements, respectively. The pattern observed with LFUSI tau seed (Figure 5C) is composed of strong negative correlations with glucose metabolism in the lateral inferior-temporal gyri, fusiform gyri, angular gyri, and precuneus, which resembles the significant regions obtained with the SVD-based tau scores (i.e., Figure 2B). As observed in Figure 5D, tau in the RANG seed was weakly related to glucose metabolism and the strongest negative correlations were observed only within the right angular gyrus itself and the posterior cingulate cortex. Notice also that the local tau in the RANG seed was associated with an increase of glucose metabolism in areas spread all over the cortex. Notably, the inclusion of SVD-based β-amyloid measures in the model (i.e., model (2) with *XSeed*_Tau_ instead of *X*_Tau_) decreases the strength and spatial extent of the local-to-distributed patterns of the tau-glucose metabolism relationship (see Figure 6). In fact, the stronger and more spatially extensive negative correlations between tau and glucose metabolism after removal of the distributed β-amyloid scores corresponded to the LFUSI seed (Figures 6A), and are only particularly strong within the left fusiform gyrus itself and the left lateral inferior temporal gyrus. On the other hand, the (weak) negative correlations for the RANG case appear restricted to the left lateral inferior temporal-parietal regions, while much larger areas of positive correlations were enhanced throughout the brain (Figures 6B).

**Figure 5.**
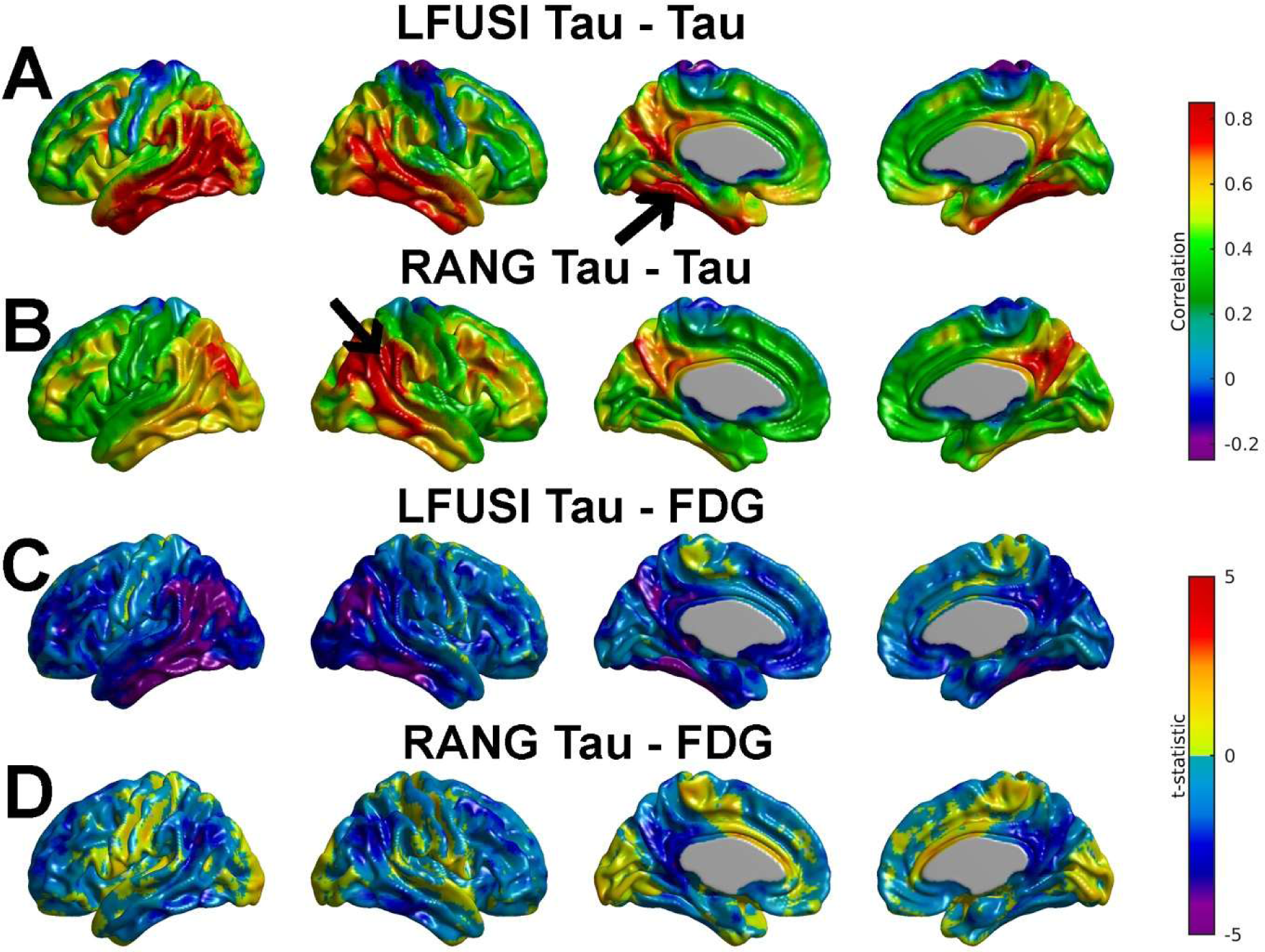
(A, B) Spatial distribution of the tau network associated with the left fusiform and the right angular seeds. (A)The seed-based tau network relative to the LFUSI seed appears to be strong and spatially extended beyond the fusiform gyrus and covers large bilateral areas of the temporal-parietal lobes and precuneus. (B) The RANG tau network is particularly strong within the right angular gyrus itself and within the bilateral precuneus and posterior cingulate cortex. (C, D) Assessment of the LFUSI and RANG seed tau measurements on FDG PET. (C) The LFUSI tau seed shows strong negative correlations with glucose metabolism in the lateral inferior-temporal gyri, fusiform gyri, angular gyri, and precuneus. (D) The RANG tau seed was only weakly related to the reduction of glucose metabolism.

**Figure 6.**
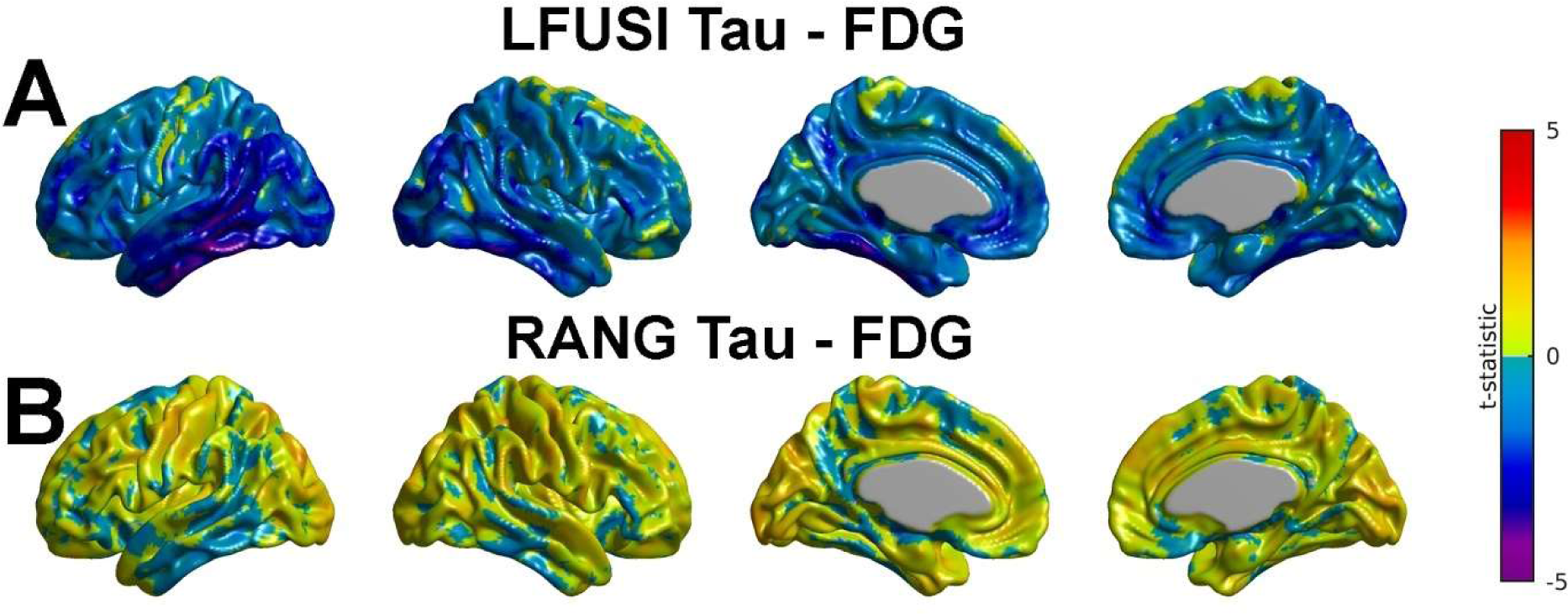
Assessment of LFUSI and RANG tau on FDG after adjustment of β-amyloid. (A) The LFUSI tau seed shows spatially extended negative correlations between tau and glucose metabolism particularly within the left fusiform gyrus itself and the left lateral inferior temporal gyrus. (B) The negative correlations for the RANG tau seed appear restricted to the left lateral inferior temporal-parietal regions. Large areas of increase in glucose metabolism are associated with RANG tau after adjustment for the β-amyloid effect.

**Table 2.** Local correlations between tau and FDG for different seed locations.

| **Seed** | **Tau SUVR** | **FDG SUVR** | **Correlation** |
| --- | --- | --- | --- |
| Left ENT | 1.28 ± 0.25 | 1.15 ± 0.11 | -0.35 |
| Right ENT | 1.27 ± 0.25 | 1.16 ± 0.11 | -0.38 |
| Left INT | 1.35 ± 0.34 | 1.50 ± 0.18 | -0.45 |
| Right INT | 1.22 ± 0.33 | 1.40 ± 0.20 | -0.27 |
| Left PCC | 1.25 ± 0.24 | 1.85 ± 0.24 | -0.27 |
| Right PCC | 1.24 ± 0.23 | 1.86 ± 0.24 | -0.27 |
| Left ANG | 1.14 ± 0.28 | 1.42 ± 0.21 | -0.28 |
| Right ANG | 1.13 ± 0.24 | 1.43 ± 0.23 | -0.21 |
| Left FUSI | 1.28 ± 0.26 | 1.45 ± 0.13 | -0.44 |
| Right FUSI | 1.29 ± 0.29 | 1.45 ± 0.14 | -0.42 |

## Discussion

We have proposed an SVD-based voxelwise cross-correlation analysis to reveal whole brain associations between FDG and tau PET images from the ADNI study. Our analysis also assessed the impact of widespread, spatially distributed β-amyloid on the relationship between glucose metabolism and tau. Our analysis highlights that the set of spatially distributed regions implicated with the reduction of glucose metabolism are the areas typically demonstrating abnormal tau uptake during the initial stages of AD (e.g., medial temporal lobe and lateral inferior temporal gyri). The individual SVD-based tau scores showed a significant association with glucose metabolism independently of the global levels of β-amyloid burden. Further, the relationship between the SVD-based scores of spatially distributed tau and glucose metabolism still holds after accounting for the remote effect of spatially distributed β-amyloid. In contrast, our seed-based correlation analysis indicated that β-amyloid has a greater impact on the association between spatially local tau and glucose metabolism. Taken together, our findings suggest that widespread β-amyloid facilitates the relationship between local tau and the reduction of glucose metabolism, but it has less influence over the effect of more spatially distributed tau beyond the medial temporal lobe.

Our SVD-based tau scores maximally correlated to FDG showed a spatial representation mainly characterized by positive high loading values within the entorhinal cortex and the lateral inferior temporal gyri, regions which Ossenkoppele *et al.* and Cho *et al.* [41,42] demonstrated to consistently show prominent tau uptake in the initial stage of AD. Note that having abnormally elevated tau uptake levels is a condition that does not automatically guarantee a strong association with FDG. Indeed, while tau uptake levels are usually assessed with first-order statistics (e.g., individual or average PET maps), their association with any other PET modality, such as FDG, must be evaluated with second-order statistic (e.g., cross-correlation analysis via SVD), and it is well known that first- and second-order statistics do not need to necessarily correlate with each other. A clear example of such a potential discordance between uptake and cross-correlation can be found from Carbonell *et al.* [14] in the case of β-amyloid and its relationship with FDG. Indeed, our seed-based correlation analysis in Carbonell *et al.* [14] showed that regions with relatively low β-amyloid burden, like the fusiform gyrus, exhibited a much stronger association with a decreasing glucose metabolism than other regions with a higher β-amyloid load. Thus, the fact that the spatial loadings associated to our tau scores follow the same spatial patterns of tau uptake suggests that the initial processes of tau seeding within the entorhinal cortex and spreading to the inferior temporal cortex have an immediate influence on the reduction of glucose metabolism. In contrast, measures of distributed β-amyloid have a weaker impact on glucose metabolism, and it is only noticeable when the associated spatial loadings are widely distributed all over the cortex, indicating a more advanced stage of pathological signs of AD.

While our SVD approach was useful for deriving maximally correlated tau and FDG scores, a more confirmatory analysis using GLM was necessary to assess the voxelwise effect of those tau scores and the APOE ε4 genotype on the FDG PET images. That modelling approach showed that, in agreement with Strom *et al.* [31], there was no statistically significant effect of the APOE ε4 genotype on glucose metabolism after adjusting for tau. This finding also goes in the same direction as our more recent study, where we demonstrated that spatially distributed β-amyloid, rather than the APOE ε4 genotype, was related to the reduction of glucose metabolism in MCI [14]. Given that Sepulcre *et al.* [43] found the APOE ε4 genotype to play a central role in tau propagation patterns, we could envision a mechanism where the genetic profile of individuals with MCI has only an indirect effect on glucose metabolism. Thus, it seems that the main contribution of APOE ε4 can be related to tau propagation, which ultimately has a much more direct impact on the reduction of glucose metabolism. More importantly, our results showed that the SVD-based tau scores were statistically significantly correlated with the reduction of glucose metabolism in extended areas of the cortex after adjusting for the indirect effect of the APOE ε4 genotype. As a basis for comparison, equivalent SVD-based scores of distributed β-amyloid were only weakly related to glucose metabolism and restricted to focal areas of the entorhinal cortex, lateral inferior temporal gyri, and angular gyri. It is worth noting that the previously mentioned regions are precisely those AD-vulnerable areas where tau had the strongest impact on glucose metabolism. Correspondingly, it is not surprising that such a small set of regions be associated with a reduction in glucose metabolism for both amyloid and tau scores since our recent analysis in Carbonell *et al.* [34] highlighted that, in those regions, widely spread β-amyloid was maximally correlated to tau. Thus, the reduction of glucose metabolism in those anatomical areas seems to be primarily driven by tau. In the current study, this hypothesis was in fact corroborated by the inclusion of the interaction term *Amyloid x Tau* in our modeling, where the weak association between β-amyloid and metabolism turned out not to be statistically significant after adjustment for the tau effect. Thus, our results are in correspondence with the causal mediation analysis presented by Bilgel *et al.* [11], where it was shown that β-amyloid has only an indirect effect on glucose metabolism and pointed to tau in the entorhinal cortex and the inferior temporal gyri as the main drivers of neurodegeneration.

In contrast to previous studies from Adams *et al.* and Rubinski *et al.* [32,33] that characterized the tau-metabolism relationship using a dichotomous global status (e.g., positive and negative) of β-amyloid levels, we followed a continuous modelling approach for the distributed scores of β-amyloid. As pointed out by Carbonell *et al.* [12], the dichotomization of β-amyloid measurements lowers the power to detect true statistical effects and thereby limits our capacity to understand the “continuum” effect of different levels of β-amyloid on the tau-glucose metabolism relationship. The inclusion of the interaction term *Amyloid x Tau* in our formulation did not only allow us to assess the synergistic effect of tau and β-amyloid on glucose metabolism, but also provided a tool to estimate the strength of the tau-metabolism association at any level of β-amyloid in a continuum. Our results showed that the gradual deposition of β-amyloid appears to modulate the spatial association between distributed measures of tau and glucose metabolism. Such association is spatially characterized by a continuous reduction of glucose metabolism in areas that follow a remarkable resemblance to the proposed topography of Braak’s staging [44–46] of tau spreading from the medial temporal lobe (stage I/II) to nearby lateral temporal regions and posterior cingulate cortex (stage III/IV) to finally reach neocortical regions (stage V/VI). Indeed, the strongest pattern of tau-related reduction in glucose metabolism corresponded to very high levels of β-amyloid, which follows the line of similar findings for cognitively normal subjects [27,32] and AD patients [23,26,31] with positive β-amyloid status.

Notice that our “amyloid-modulation” analysis also revealed small clusters (e.g., pars opercularis) where higher tau was (weakly) associated with increases in glucose metabolism. However, those small clusters tend to reduce their spatial extent and strength with the gradual increase of β-amyloid. Similar positive associations have already been reported in previous studies involving cognitively normal subjects by Hanseeuw *et al.* and Adams *et al.* [27,32]. For instance, Hanseeuw *et al.* [27] demonstrated higher levels of tau in the inferior temporal region to be locally associated with increased glucose metabolism in CN subjects. Also in CN subjects, Adams *et al.* [32] showed that pairwise correlations between ROI measures of tau and FDG were predominantly positive for a sample of negative amyloid subjects. Similarly, Rubinski *et al.* [33] indicated that, for MCI subjects, local increases in tau are associated with local increases in glucose metabolism at low levels of β-amyloid. In this regard, our results showed that, although not statistically significant, local increases of tau within the right angular gyrus were associated with increases of glucose metabolism within areas of the occipital cortex, the paracentral lobules, and the anterior cingulate cortex. Further, those positive sign associations were extended to large areas of the cortex after adjustments for the effect of distributed β-amyloid. It is a remarkably finding since the angular gyri turned out to be regions of prominent reduction in glucose metabolism associated with distributed tau. In combination, these findings reinforce the idea that a tau measure taken from those areas typically associated with initial seeding and spreading of tau pathology would have the greatest role in the reduction of glucose levels in remote neocortical regions that are supposed to be impacted by tau at a later stage. Henceforth, the tau-related increases in glucose metabolism in areas such as the angular gyri have been typically interpreted, for example by Hanseeuw *et al.* [27], as a compensatory mechanism of increased neuronal activity driven by the impact of tau spreading. Even though the molecular mechanisms causing the tau-related hypermetabolism at low β-amyloid levels are still largely unknown, Adams *et al.* [32] suggested that it might possibly represent a normal aging process. Alternatively, Rubinski *et al.* [33] proposed a scenario where tau enhances neuronal hyperactivity at lower levels of β-amyloid, but reduces neuronal function with the gradual accumulation of β-amyloid.

In any case, a plausible mechanistic explanation for the uncovered tau-metabolism associations, as well as their modulation by the concurrent action of β-amyloid would necessarily involve multi-factorial, integrative scenarios of the AD progression. From a purely brain connectivity perspective, one must consider the relationship between tau deposition, abnormal structural and functional connectivity, and underlying amyloid-dependent patterns of spread [47–50]. For instance, modelling approaches by Vogel *et al.* [51] support the notion that tau pathology propagates through neuronal connections, likely with seed origin in the medial temporal cortex, a process that is accelerated under the influence of a gradual increase in β-amyloid accumulation.

On the other hand, given the well-known strong association among multiple neuroanatomical, neurophysiological, and neurovascular systems as per Iturria-Medina *et al.* [52] in neurodegenerative disease progression, it is unlikely that pathologic alterations caused by toxic β-amyloid and/or tau proteins would not alter the brain’s metabolic equilibrium or vice-versa. For instance, according to Qosa *et al.* [53], disruptions in the neurovascular integrity and neurovascular coupling can affect misfolded proteins clearance, leading to high intraparenchymal β-amyloid and tau protein concentration levels. Correspondingly, Iadecola *et al.* [54] demonstrated that gradual increases in β-amyloid and tau may induce brain-blood barrier (BBB) alterations, thereby damaging endothelial cells, pericytes, vesicular transport, and calcium balance. Hence, the combined effect of the previously mentioned processes would lead to an inevitable alteration in the brain metabolism, producing a multi-factorial and continuous degenerative cycle.

### Conclusions

We have explored whole brain associations between glucose metabolism, tau, and APOE ε4 genotype in a sample of MCI subjects. Our analysis also assessed the impact of β-amyloid on the spatial relationship between glucose metabolism and tau. By exploring the large-scale, cross-correlation between tau and FDG PET images with the SVD approach, our analysis revealed key findings, including: (1) reduction of glucose metabolism is not associated with APOE ε4 genotype, (2) spatially distributed tau has a much greater role in the reduction of glucose metabolism than β-amyloid, (3) tau-related reduction of glucose metabolism was pronounced even after adjustment for β-amyloid, (4) gradual increase in β-amyloid continuously modulates the tau-related reduction in glucose metabolism, and (5) local tau in initial seeding regions had a prominent role the reduction of glucose levels in remote neocortical regions. To our knowledge, this study is the first to relate glucose metabolism with tau and β-amyloid from a whole brain network perspective that accounts for distributed-to-distributed and local-to-distributed patterns of cross-correlation.

## Acknowledgments

Data collection and sharing for this project was funded by the Alzheimer’s Disease Neuroimaging Initiative (ADNI) (National Institutes of Health Grant U01 AG024904) and DOD ADNI (Department of Defense award number W81XWH-12-2-0012). ADNI is funded by the National Institute on Aging, the National Institute of Biomedical Imaging and Bioengineering, and through generous contributions from the following: AbbVie, Alzheimer’s Association; Alzheimer’s Drug Discovery Foundation; Araclon Biotech; BioClinica, Inc.; Biogen; Bristol-Myers Squibb Company; CereSpir, Inc.; Cogstate; Eisai Inc.; Elan Pharmaceuticals, Inc.; Eli Lilly and Company; EuroImmun; F. Hoffmann-La Roche Ltd and its affiliated company Genentech, Inc.; Fujirebio; GE Healthcare; IXICO Ltd.; Janssen Alzheimer Immunotherapy Research & Development, LLC.; Johnson & Johnson Pharmaceutical Research & Development LLC.; Lumosity; Lundbeck; Merck & Co., Inc.; Meso Scale Diagnostics, LLC.; NeuroRx Research; Neurotrack Technologies; Novartis Pharmaceuticals Corporation; Pfizer Inc.; Piramal Imaging; Servier; Takeda Pharmaceutical Company; and Transition Therapeutics. The Canadian Institutes of Health Research is providing funds to support ADNI clinical sites in Canada. Private sector contributions are facilitated by the Foundation for the National Institutes of Health (www.fnih.org). The grantee organization is the Northern California Institute for Research and Education, and the study is coordinated by the Alzheimer’s Therapeutic Research Institute at the University of Southern California. ADNI data are disseminated by the Laboratory for Neuro Imaging at the University of Southern California.

## Author contributions

FC and BJB designed research; FC, CM, APZ, and BJB performed research; FC and BJB analyzed the data and drafted the paper.

## Compliance with ethical standards

### Ethical approval

All procedures performed in studies involving human participants were in accordance with the ethical standards of the institutional and/or national research committee and with the 1964 Helsinki declaration and its later amendments or comparable ethical standards.

### Informed consent

Informed consent was obtained from all individual participants included in the study.

---

**Supplementary Figure 1.**
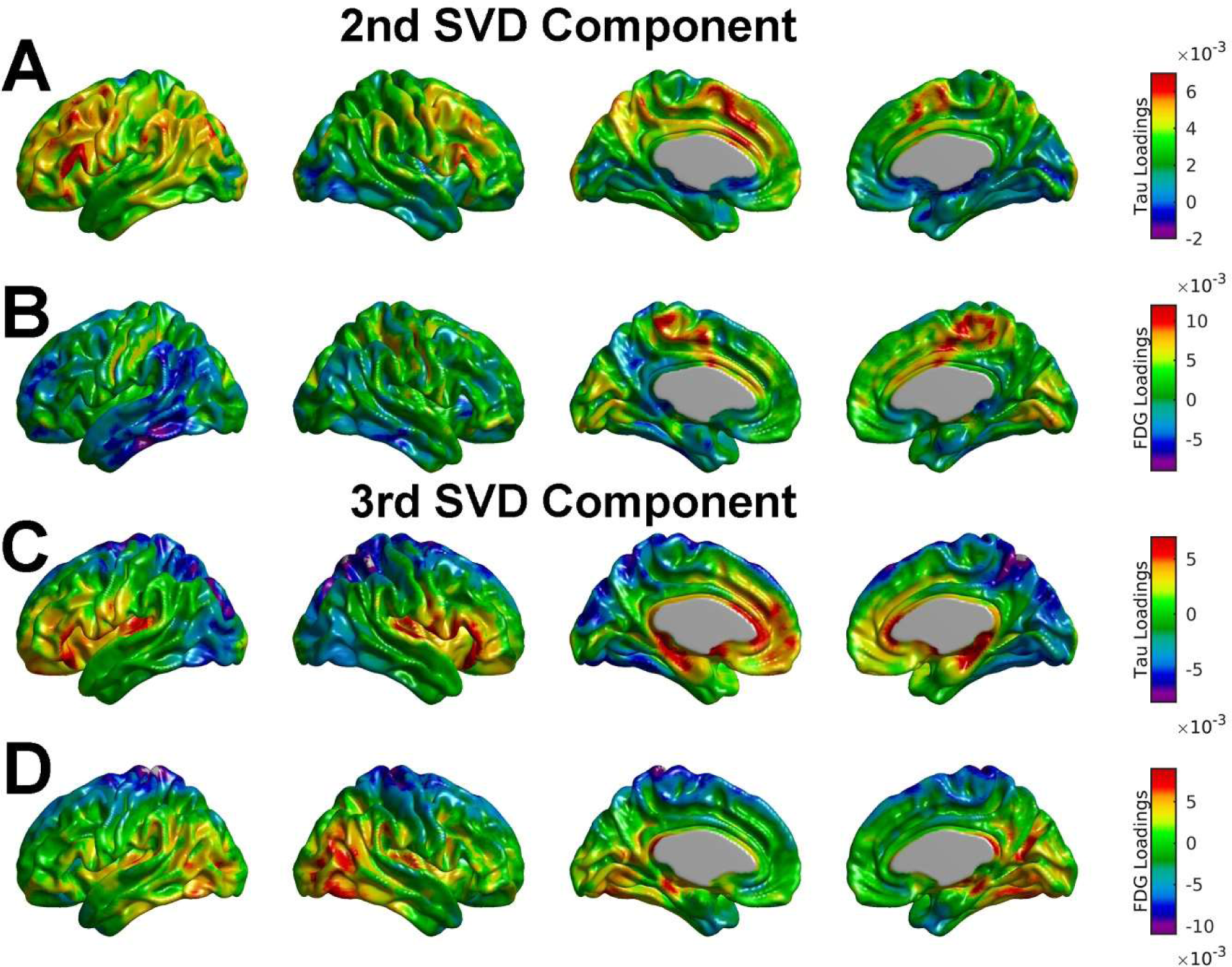
Spatial loadings for tau and FDG corresponding to the second and third SVD components. (A, B) The second component highlights the situation where an increase in tau within areas of the lateral and medial frontal lobules is associated with both a reduction of glucose metabolism in the lateral inferior temporal gyri and an increase of glucose metabolism within the medial frontal-parietal cortex. (C, D) The third SVD component reflects that the increase of tau within the medial orbital-frontal cortex and the parahippocampal gyri is associated with an increase of glucose metabolism within the temporal-occipital cortex.

**Supplementary Figure 2.**
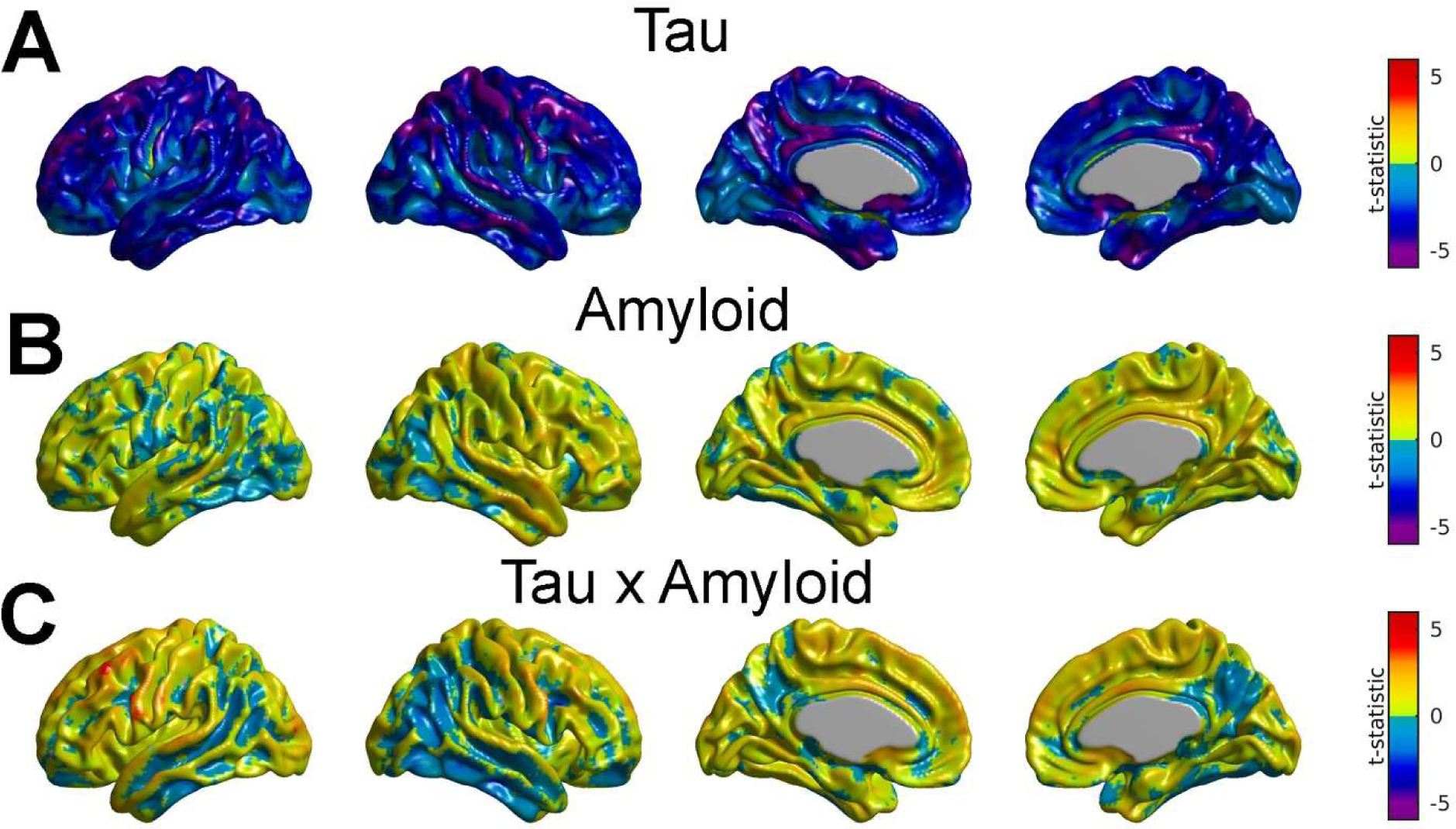
Statistical assessment of the alternative amyloid scores resulting from the SVD analysis between β-amyloid and tau PET images on FDG PET. (A) Extended areas of reduction in glucose metabolism associated with tau scores after controlling for the effects of the alternative β-amyloid scores. (B, C) Neither the main effect of β-amyloid nor the interaction between β-amyloid and tau produced statistically significant results.

**Supplementary Figure 3.**
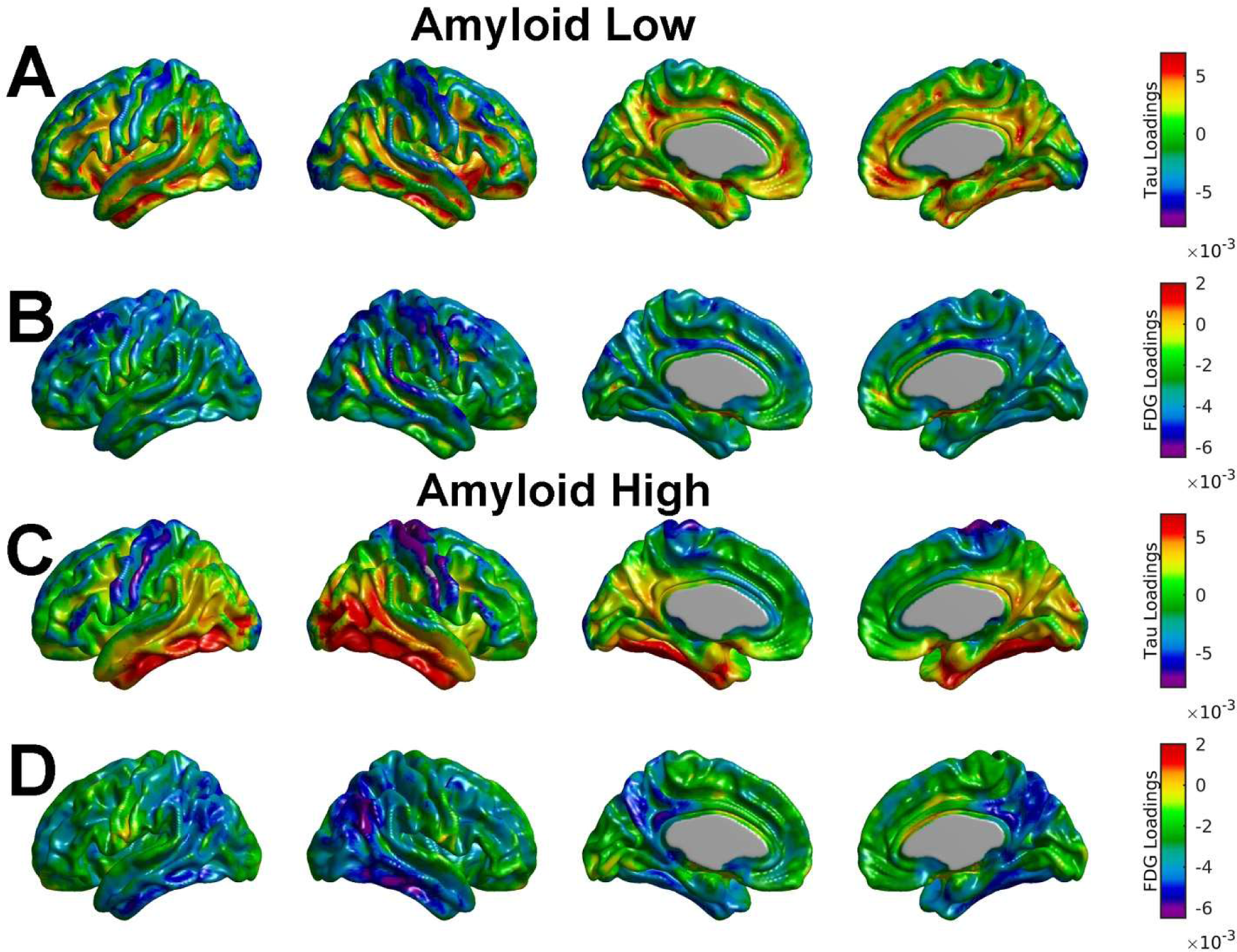
SVD analysis for the two independent groups corresponding to the AβH and AβL subjects. The tau loadings appear to be stronger for the Aβ_H_ case, particularly for the lateral inferior temporal gyri and the fusiform gyri. Also, the negative FDG loadings are only strong for the Aβ_H_ case.

